# Ticks Home in on Body Heat: A New Understanding of Ectoparasite Host-Seeking and Repellent Action

**DOI:** 10.1101/564179

**Authors:** Ann L. Carr, Vincent L. Salgado

## Abstract

Ticks are second only to mosquitoes as vectors of disease to humans and animals. Commercial insect repellents reduce or prevent potentially infectious tick bites by disrupting tick host-seeking behavior. Tick host-seeking is mainly ascribed to the Haller’s organ, a complex sensory structure on the tick foreleg that detects odors, carbon dioxide and heat ^4–7^, but these host-seeking mechanisms and the mechanism of their disruption by repellents are not well understood^1,2^. There is anecdotal evidence that ticks and other ectoparasites are attracted to heat, but it has never been demonstrated that they use radiant heat to detect hosts at a distance. In fact, previous attempts to do this have concluded that radiant heat is not used by ticks. Here we show that *Amblyomma americanum* and *Dermacentor variabilis* ticks can sense and home in on a human from several meters away, guided by radiant heat sensed by the Haller’s organ, and specifically the capsule, a covered spherical pit organ. An aperture in the capsule cover confers directionality and highly reflective interior surfaces of the capsule provide high sensitivity. Low concentrations of the insect repellents DEET, picaridin, 2-undecanone, citronellal and nootkatone eliminate thermotaxis without affecting olfactory-stimulated host-seeking behavior. Our results demonstrate that the tick Haller’s organ capsule is a radiant heat sensor used in host-finding and that repellents disrupt this sense at concentrations that do not disrupt olfaction. We anticipate this discovery to significantly aid insect repellent research and development of innovative strategies for protection against ectoparasites and vector-borne disease.

## Main Text

Ticks transmit the widest array of pathogens of any arthropod vector, including viruses, bacteria, fungi, protozoa and helminths, and are second in importance only to mosquitoes as vectors of disease to humans and animals^1^. The American dog tick *Dermacentor variabilis* (Acari: Ixodidae) and the lone star tick *Amblyomma americanum* (Acari: Ixodidae) are the two most abundant ticks in the US^2^; both have the capacity to transmit ehrlichiosis, tularemia and Rocky Mountain spotted fever. Also of public health concern is the increasingly prevalent galactose-alpha-1,3-galactose red meat allergy associated with *A. americanum* bites ^2,3^.

Ticks find hosts using the Haller’s organ on the foreleg to detect odors, carbon dioxide and heat ^4–7^, but the only study to investigate the role of heat in tick host-seeking concluded that radiant heat is not used ^4^. The part of the Haller’s organ known as the capsule is thought to serve an olfactory function ^4,8–10^ but its camera-lucida-like structure as a covered, sensilla-containing spherical pit with an aperture ^8,11^ seems better suited to detect radiation. Although the aperture would severely hinder odorant diffusion, it would confer directionality for radiation-sensing by restricting field of view. We show that previous studies on heat sensing by ticks were flawed and that the Haller’s organ capsule is indeed an exquisitely sensitive radiant heat sensor that guides host-finding. We further show that exposure to common tick repellents disrupts this thermotaxic behavior, which we propose to be a novel mode of action of repellents.

Questing unfed virgin adult *A. americanum* females placed on the floor in the center of a 10 cm wide X 10 cm high X 1m long thermotaxis arena (Fig. 1) were strongly attracted to a warm surface (target) at one end of the arena, 50 cm away, when it was at 37 °C or warmer. Most ticks moved toward the warm target almost immediately, and of the few that initially moved toward the opposite (cold) end of the arena, most reversed direction, so that after 5 min only one tick of 48 remained within 10 cm of the cold wall, while 40 were within 10 cm of the 37 °C target. Seven ticks ended the trial between these end regions (Fig. S1, Movie S1). With the target at 22 or 30 °C, many ticks did not move far from the origin, and an approximately equal number moved toward either end, with a strong tendency to continue moving in the initial direction. Results were similar for *A. americanum* males (Fig. S1) and both sexes of *D. variabilis* (Fig. S2).

**Figure 1.**
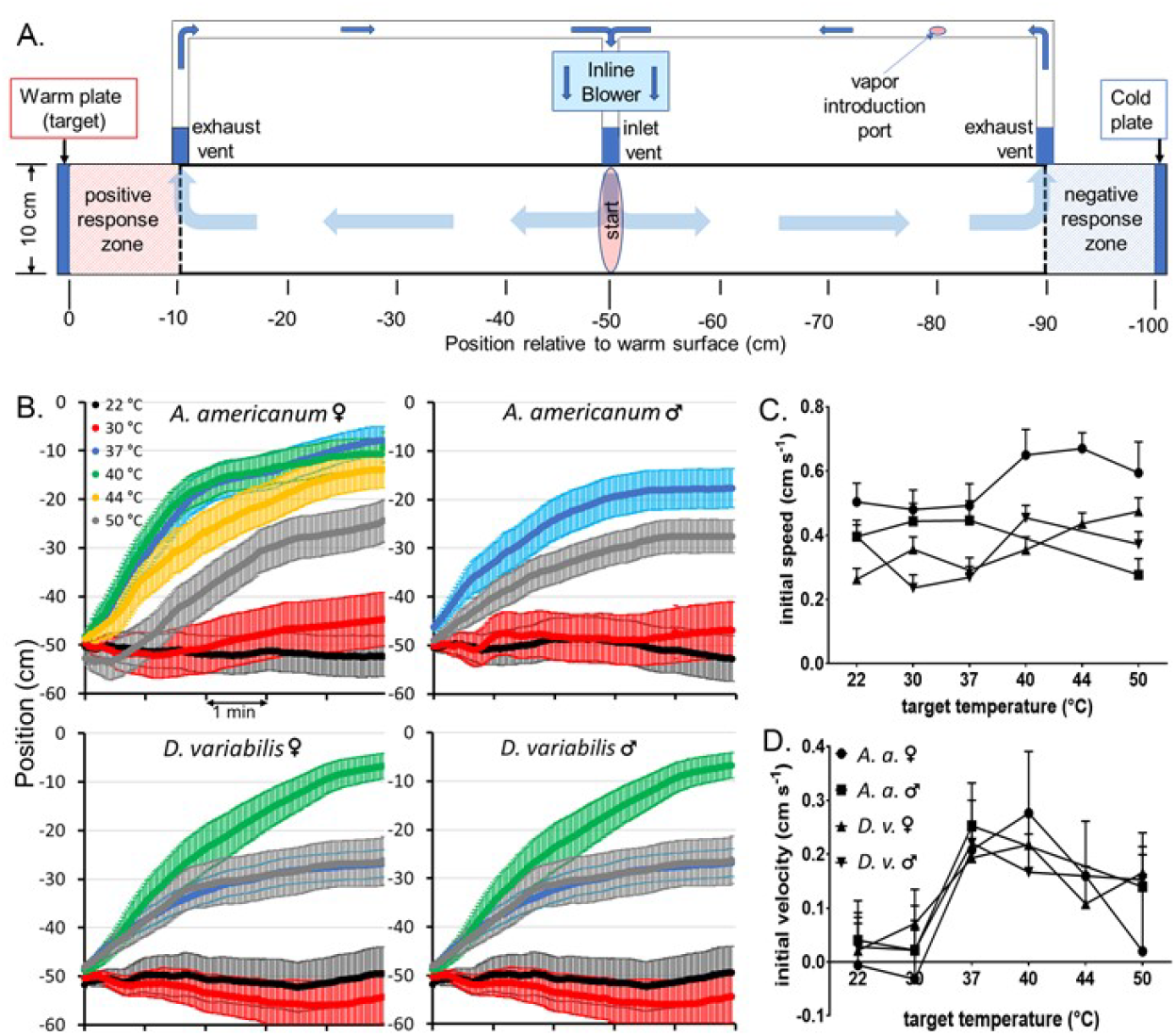
A. Thermotaxis arena with closed air circulation system ^12^. B. Longitudinal position vs. time in the thermotaxis arena of the indicated ticks, after placement in the center at −50 cm with respect to the warm target at 0 cm. Horizontal axis units are minutes. C. Initial speed, and D. velocity of ticks during the first 10 seconds after release.

Net movement and average initial velocity toward the target began immediately when it was at 37, 40 or 44 °C, and except in the case of *A. americanum* females, also at 50 °C (Fig. 1B, D), indicating that ticks easily detected the warm target from 50 cm away. Initial speed did not vary consistently with temperature (Fig. 1C), indicating that temperature guides but does not stimulate movement. Initial velocities declined at target temperatures above 40 °C (Fig. 1D).

All ticks studied showed a significant warm/cold preference at 37 °C and above for final choice and initial movement (Fig. 2A, B). The fraction choosing an end region was always higher for initial movement compared to final choice, because some ticks stopped moving before reaching either end. A preference index (PI) ranging from −1 to +1, with 0 indicating no preference, quantifies warm preference (Fig. 2C, D). PI was near zero at 22 and 30 °C. Initial PI at 37 °C was between 0.7 and 0.95, indicating that at least 70 % of ticks detect the 37 °C target at 50 cm. Except for *A. americanum* males, initial PI showed a sharper optimum than final PI, indicating that temperatures above 40 °C may be initially inhibitory.

**Figure 2.**
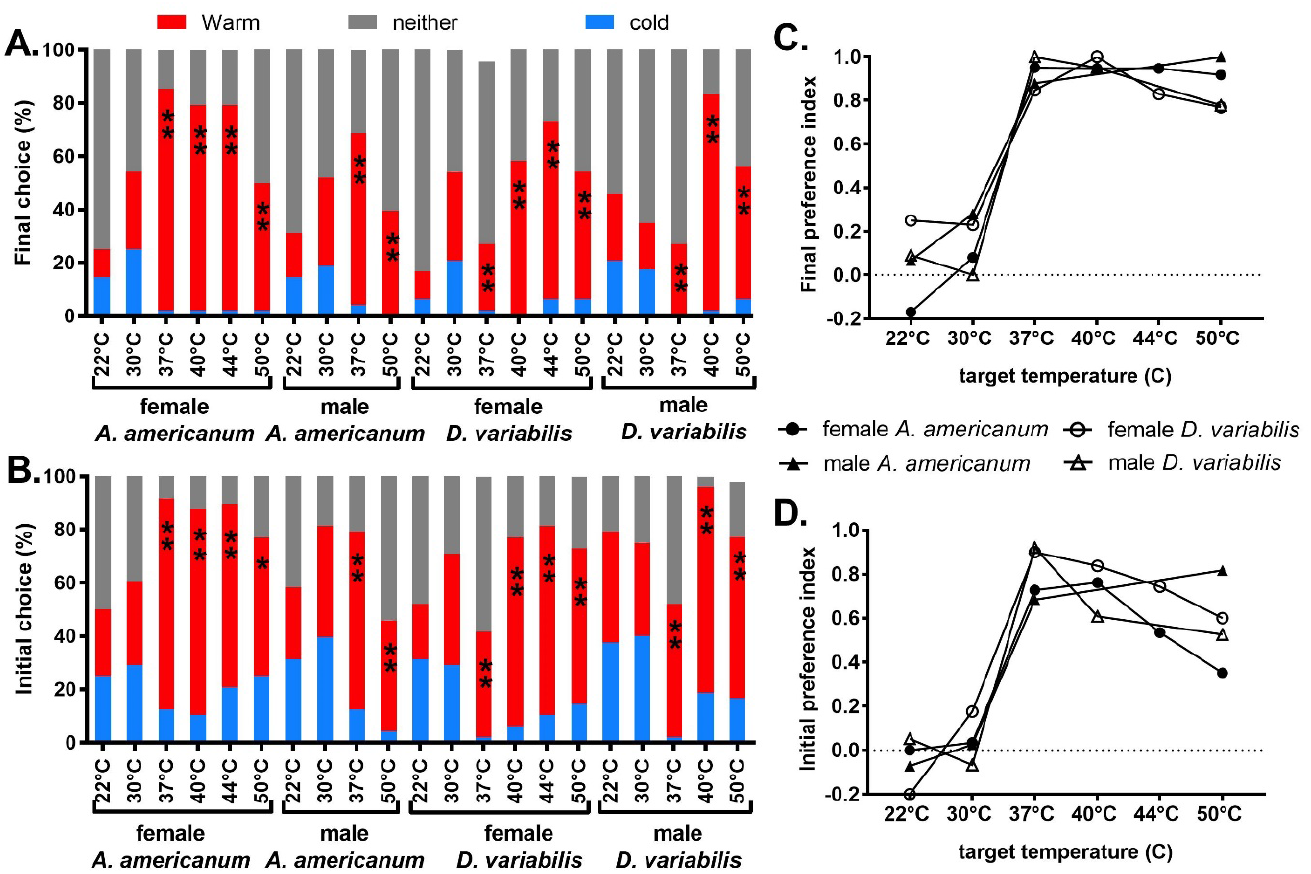
Responses of ticks in thermotaxis trials of Figs. 1, S1 and S2 scored as choice assays. (A) Final choice of ticks at the end of a 5 min trial scored as “warm” or “cold” if within 10 cm of the corresponding end, or “neither” if between these end areas. (B) Initial choice scores based on direction of the first movement of more than 15 cm if the tick eventually moved at least 25 cm from the origin. Ticks that failed to move at least 25 cm were scored “neither”. (C) Final warm preference index (PI), defined as (warm-cold)/(warm+cold) ^20^. (D) Initial warm PI. Significance of warm/cold difference: **: p < 0.001; *: p < 0.05.

To better assess tick heat-sensing range, we started *A. americanum* females at the far end of the arena, 100 cm from the target. With the target at 22 °C, many ticks immediately began moving toward it, ending near the center of the arena, at −55 ± 4 cm (mean ± SEM; Fig.3A). With the target at 40 °C to radiate energy at the same rate as human skin at 37 °C ^12^, average movement toward it was much faster from the start and the ticks reached an average distance of only 16.5 ± 4.5 cm from it (Fig. 3A). From the distribution of initial velocities (Fig. 3B), 45 of the 88 ticks showed initial movement toward the target when at 22 °C, and can be considered to be moving spontaneously, whereas the remaining 43 were not. With the target at 40 °C, 76 ticks began moving toward it immediately, representing 45 that were moving spontaneously and 31 that were thermotaxing. Since 31 of the 43 ticks that were not moving spontaneously, or 72%, began thermotaxing immediately from 1m away, we can conclude that at least 72% of adult *A. americanum* females detect the 10 × 10 cm target at a distance of 1 m, just as many as at 0.5 m. A longer arena or smaller target is needed to assess the range of tick heat detection, but we can conclude that since an adult human torso is approximately 4 times the width of our target, a tick could detect an adult human at a distance of < 4 meters under ideal conditions.

**Figure 3.**
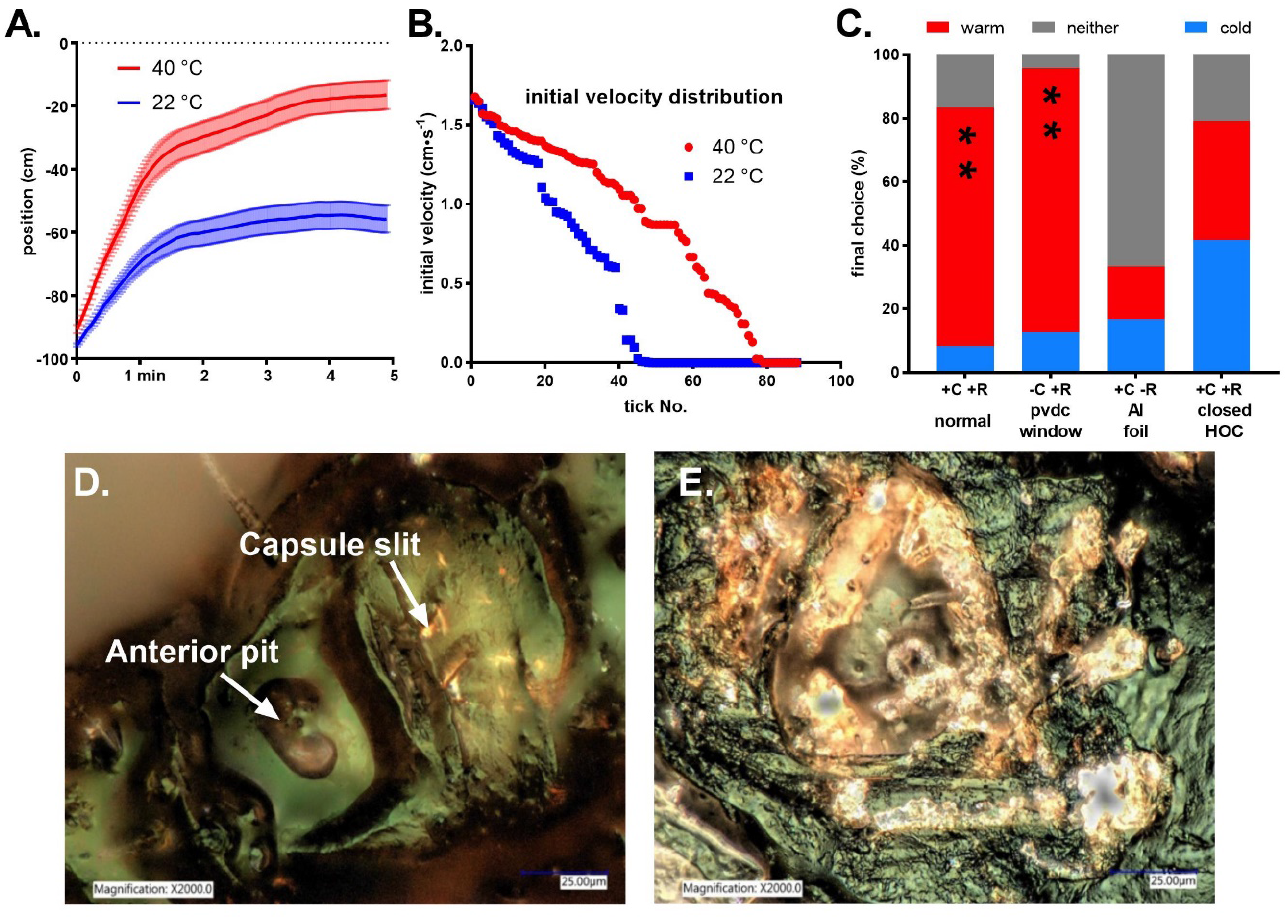
Sensitivity and mechanism of tick heat sensing. A. Mean longitudinal position (± SEM) over time (s) of 88 *A. americanum* females (measured in 11 groups of 8) after placement adjacent to the cold end wall at −100 cm with respect to the target wall that was set to either 22 or 40 °C. B. Distribution of initial velocities of all 88 ticks tested at 22 or 40 °C, averaged over the first 10 s, from fastest to slowest. C. Final choice scores in thermotaxis trials of *A. americanum* females placed in the center of the arena, with the target at 40 °C, under normal conditions, where convection and radiation from target to ticks are both possible (+C +R, normal), with an IR-transparent pvdc food wrap barrier 5 cm from each end wall to block any possible convective air currents (-C +R, pvdc window), with both end walls clad with aluminum foil to eliminate radiant heat emission and leave only the possibility of heat transfer by convection (+C -R, Al foil), and finally in the normal arena but with both Haller’s organ capsule apertures closed with paraffin wax (+C +R, closed HOC). (** = p < 0.001 for warm/cold difference). D. Female *A. americanum* Haller’s organ viewed with epi-illumination shows hints of reflected gold light through the capsule aperture. E. Interior of the Haller’s organ capsule and cover debris appear highly reflective after cover removal with an ultrasonic micro-etcher ^12^.

The fastest walking speed of *A. americanum* females in Figure 3B was 1.65 cm•s^−1^, equal to the speed observed for *Hyalomma* ticks hunting in the field ^5^. The fastest speeds we observed for *A. americanum* males, *and D. variabilis* females and males were 1.5, 1.2 and 1.1 cm•s^−1,^ respectively.

We propose that the tick Haller’s organ capsule is a radiant heat sensor. Conduction of heat along the floor and sides of the arena was insignificant ^12^ and elimination of the possibility of convective heat transfer from the target to the ticks by interposing an IR-transparent plastic barrier had no significant impact on the percentage of ticks choosing the 40 °C target (Fig. 3C). On the other hand, when both arena end walls were clad with aluminum foil to eliminate radiant heat emission without affecting convection, or when the Haller’s organ capsule apertures were waxed over, the ticks no longer preferred the 40 °C target (Fig. 3C), confirming that the capsule is a radiant heat sensor that guides tick thermotaxis.

We reasoned that for the capsule sensillae to sense temperature of a distant source, the capsule walls and the interior surface of the cover must be reflective. Viewed with epi-illumination, the aperture of the intact capsule appeared to emit reflected light (Fig. 3D). When the cover was removed, the walls of the capsule and the inside faces of cover fragments scattered on the surrounding cuticle appeared highly reflective (Fig. 3E).

Commercial insect repellents dose-dependently reduced tick thermotaxis (Figs. 4A, B, S3). Exposure to 50 ng•cm^−3^ DEET, 54 ng•cm^−3^ picaridin and 42 ng•cm^−3^ 2-undecanone completely and reversibly abolished *A. americanum* female thermotaxis, with 50 % inhibitory concentrations of 26, 31 and 17 ng•cm^−3^, respectively (Fig. S3B). Thermotaxis was also disrupted by citronellal and nootkatone (Fig. 4B), but fewer trials were conducted because their residues were difficult to remove from the arena. Individual ticks exposed to any of the repellents behaved as if there were no warm target, beginning to move immediately and continuing in the direction in which they started, but showing no clear end wall preference (Fig. S3A, Movie S1). Number of ticks making a choice (Fig. S3B) and initial speed (Fig. S3C) did not decline with increasing repellent concentration, indicating that repellents affect heat detection but not host-seeking behavior. We next examined the effect of DEET on the olfactory response to CO_2_, a necessary and sufficient stimulant of host-seeking behavior ^6,13^. In control Y-tube bioassays testing medical grade air in both arms (Fig. S4), 32% of ticks made a choice, with approximately equal numbers going to either arm (Fig. 4C, sham CO_2_). Introducing 4% CO_2_ into one arm, the concentration in human breath, resulted in 52 % of ticks making a choice - primarily the arm with CO_2_. Exposure to 50 ng•cm^−3^ DEET had no effect on tick detection of CO_2_ in Y-tube bioassays (Fig. 4C), confirming that DEET at this concentration disrupts thermotaxis but not host-seeking behavior.

**Figure 4.**
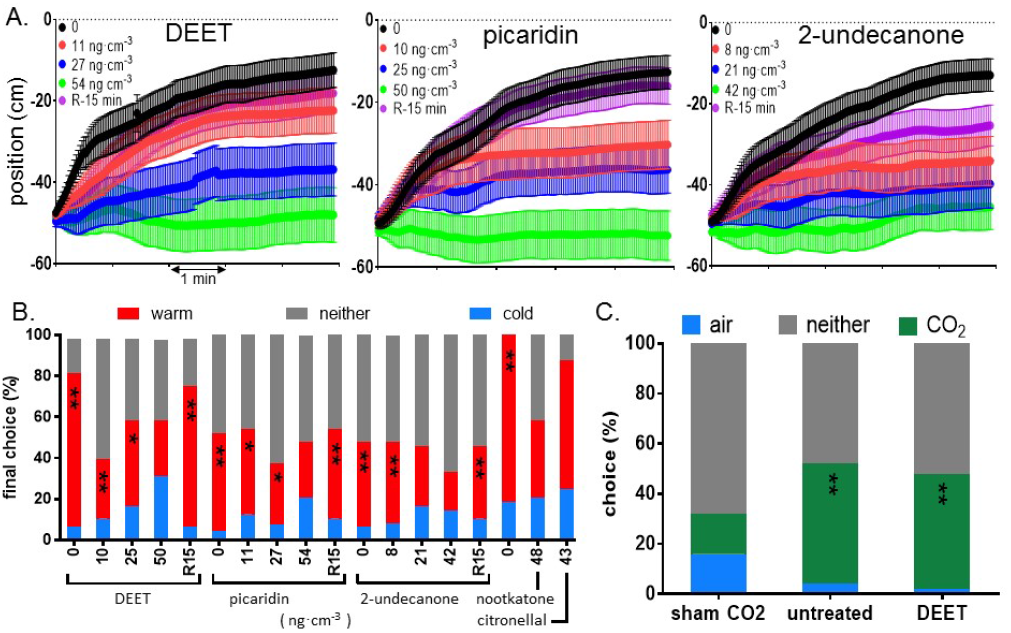
Disruption of thermotaxis by repellents. A. Mean longitudinal position (± SEM) in thermotaxis trials of unfed virgin adult *A. americanum* females exposed to the indicated repellent vapor concentrations (ng•cm^−3^) or after 15 minutes’ recovery from the highest concentration (R). Ticks were placed into the center of the arena, at −50 cm with respect to the 40 °C target wall. B. Final position after 5 minutes, scored as in Fig. 2A. C. Y-tube choice assays with unfed virgin adult *A. americanum* females under three conditions. Sham CO_2_: both tubes contained pure air. Untreated: air in one tube contained 4% CO_2_. DEET: air in one tube contained 4% CO_2_ and ticks were exposed to 50 ng•cm^−3^ DEET. Significance of warm/cold or CO_2_/air difference: **: p<0.001; *: p<0.05.

We have shown here for the first time that ticks thermotax guided by a radiant heat sense in the Haller’s organ capsule sensitive enough to locate a human body 4 meters away. Radiant body heat includes wavelengths between 5 and 20 μm ^14^, well outside the spectral sensitivity of rhodopsins, and biological radiant heat detectors are normally thermosensors arranged in a pit such that they are warmed by radiant heat from the target in preference to that from the self ^15^. We have demonstrated that the Haller’s organ capsule is in fact such a “pit organ”, with an aperture and a reflective interior lining for high directionality and sensitivity. The ability of *A. americanum* females to detect a 10 cm wide target at 1 m indicated a field of view of 6 degrees or less. Recent findings have shown that the tick Haller’s organ, but not specifically the capsule, is required for attraction to 880 nm near-infrared light ^16^. While well outside the range of thermal IR, it is likely that the tick radiant heat sensors were warmed enough by this light to stimulate thermotaxis. However, we demonstrate for the first time that the capsule is a radiant heat sensor that plays a critical role in tick host-seeking.

The mechanism of action of arthropod repellents is not well understood. While they are generally thought to repel or to cause directed movements away from the source ^17^,we have shown that several of the most commonly used personal repellents specifically disrupt tick thermotaxis at concentrations that do not affect CO_2_-stimulated host-seeking behavior. Repellents have been studied extensively on mosquitoes, which can detect them by olfaction and gustation ^18,19^, but it is not clear whether this detection ability is the mechanism of repellency in the field. In light of the findings presented here, the roles of olfaction and heat sensing in repellent action must be re-evaluated.

## Acknowledgments

We thank Dan Houtz for advice on image analysis, Jackelyn Soto for help with epimicroscopy and Hongmei Jia for statistical analysis.

## Author Contributions

VLS: Conceptualization, data curation, formal analysis, funding acquisition, methodology, supervision, visualization, writing-original draft, writing – review and editing. ALC: investigation, visualization, writing-review and editing.

## Author Information

Authors declare no competing interests. Correspondence and requests for materials should be addressed to Dr. Vincent Salgado.

## Data and materials availability

All data are available in the main text, methods or supplementary materials.

## Supplementary Materials

Figures S1 to S5

Captions for Movie S1

### Other Supplementary Materials for this manuscript include the following

Movie S1

Table S1 –all choice data and statistical analyses.

## Methods

### Ticks

Unfed virgin adult female and male *Amblyomma americanum* (lone star tick) and *Dermacentor variabilis* (American dog tick), two weeks post-molt, were purchased from Ecto Services Inc. (Henderson, NC, USA). Both colonies were started with field-collected *A. americanum* and *D. variabilis* obtained from a single site in Stillwater, OK and maintained with subsequent collections from the same site. Adult female and male *A. americanum* and *D. variabilis* were maintained in separate containers at 23 ± 1 °C and 97% R.H., with a photoperiod of 10 h light: 14 h dark, including 1h-long dusk and dawn periods under red light at the beginning and end of each scotophase. A 10 h photophase was used because this was found to give increased host-seeking, feeding and longevity in studies with laboratory-reared adult *Aedes albopictus* and *A. aegypti* ^13^.

### Thermotaxis Assays

We used a thermotaxis assay to investigate the detection range, temperature dependence and repellent sensitivity of heat perception in ticks and to identify the sensory organ responsible for this sense. A warm surface rather than an electronic infrared radiation (IR) source was used as the target for thermotaxis since the radiant heat emitted by the warm surface would closely mimic the spectrum and intensity of thermal radiation emitted by host animals, allowing the determination of the temperature sensitivity of the radiant heat sense. Furthermore, it is important to use a large enough surface as a radiant heat source because, while the intensity of radiation from each point on the surface decreases as the square of distance, the surface area within the field of view of the radiant heat sensor increases by the same factor, so that the intensity of radiant heat entering the sensor is independent of distance as long as the target fills the entire field of view. The key parameters of a radiant heat sensor are therefore temperature sensitivity and field of view.

Thermotaxis assays were conducted in an arena consisting of a long box with interior dimensions of 10 cm x 10 cm x 100 cm (Fig. 1). The long walls and floor of the box were made of ¾ inch birch plywood painted with flat white latex paint, and the top was a tight-fitting removable lid made from ¼ inch thick Plexiglas to allow observation from above while blocking thermal radiation from outside the arena. Each end wall of the arena was the 10 cm x 10 cm blue anodized aluminum surface of a temperature-controlled thermoelectric cold plate (TEC plate, model TCP-50, Advanced Thermoelectric, Nashua, NH, USA), controlled by a TEC controller (model 1000-0649) and WinVue software v .4.011 from Vuemetrix (San Jose, CA, USA). All parts of the arena fit tightly enough that ticks could not escape. The arena was always at room temperature (22 ±1 °C), while the TEC plates presented temperature-controlled surfaces with an emissivity of 0.94 (measured with a FLIR E8 thermal imaging camera, FLIR Instruments Inc., Wilsonville, OR, USA)) at either end of the arena. All TEC plate temperatures were verified with a precision thermocouple thermometer (BAT-12, Physitemp Instruments Inc., Clifton, New Jersey, USA), with the thermocouple inserted into the center of the 1 cm-thick aluminum TEC plate through a hole drilled into one side and filled with heat-conducting paste. The controller’s feedback thermistor was in the same location. The thermal imaging camera was used to verify arena temperatures and to check for unintended infrared reflections. We specifically ensured that reflection of thermal radiation from the warm plate at 50 °C in the cold plate at RT was undetectable, as such reflections would confuse the ticks and decrease the thermotaxis toward the warm plate.

Thermotaxis trials were conducted during the day between the hours of 10:00-16:00 at room temperature (RT, 22 ± 1°C), with a relative humidity of 40%, under LED lighting as described below. One TEC plate, designated the cold plate or end wall, was programmed to present a RT surface, and the opposing TEC plate, designated the target, was programmed to present various temperatures between RT and 50°C. Control trials with both TEC plates set to RT were performed to document the normal behavior of ticks in the test arena and to ensure that there were no biases associated with the laboratory conditions or experimental set-up. Unless stated otherwise, thermotaxis trials consisted of six runs of eight ticks. Ticks were randomly selected from a total population of 250 and were acclimated to test conditions in vials next to the arena for 20 min prior to use. Ticks were released onto the floor at the center of the arena (−50 cm) and their movements were filmed for 5 min, unless noted otherwise. To eliminate contamination and positional bias, all interior surfaces of the arena were washed thoroughly with 70% ethanol between runs and the cold and warm walls were alternated every three replicates. Both TEC plates were always temperature-controlled, even when at ambient temperature. Robust tick response within the thermotaxis arena required that ticks be actively displaying the host-seeking behavior known as questing, which is stimulated by CO_2_ in the breath of hosts ^10,21^. Questing of tick test subjects was induced by brief agitation of the holding vial and by exposure to the breath of the experimenter. All ticks were questing prior to placement into the arena for thermotaxis bioassays.

Two 700 lumen LED floodlights with a color temperature of 4000 K (Utilitech Pro 1-Light, 16 Watt Portable Work Light; Utilitech) were trained on the center of the arena at angles of 45 degrees, at a horizontal distance of 65cm from either end. Thermotaxis trials were filmed from above at 1080i resolution with a Canon rebel T6i camera with an 18-55mm zoom lens (Canon, Tokyo, Japan). Each 5-minute-long MP4 video file from a run was transferred to a computer and converted to a stack of 150 JPEG images at 2-second intervals, which were loaded, in chronological order into ImageJ ^22^. The position of each tick along the longitudinal axis of the arena was tracked using the MTrack3 plug-in and reported every 2s in cm, with the warm plate located at 0 and the cold plate at −100 cm.

With the exception of temperature sensitivity trials, the temperature of the target was set to 40 °C, the temperature at which, according to the Stefan-Boltzmann law, a surface with an emissivity of 0.94, as measured for the blue anodized TEC plate surface, would radiate energy at the same intensity as human skin does at 37 °C, which has an emissivity of 0.98 ^14^.

Previous investigations that failed to find that ticks ^4^ or other hematophagous insects ^23^ were attracted by radiant heat used a test tube covered with aluminum foil as the non-emitting source and a similar tube with a layer of cellophane over the aluminum foil as the IR-emitting source. We were skeptical that the thin, transparent layer of cellophane would emit sufficient radiant heat, so we wrapped a section of a 1l graduated cylinder with aluminum foil and another section of the same cylinder with aluminum foil covered with cellophane, and filled the cylinder with 37 °C water. The thermal imaging camera measured the temperature of the uncovered glass as 37 °C and that of the aluminum foil section as 21 °C, because the foil emits almost no thermal IR itself, but rather reflects that from the room temperature surroundings. The cellophane over aluminum foil section, which was intended in the cited studies to provide radiant heat, appeared to the thermal imaging camera to be at 24.6 °C (Fig. S2) indicating that the cellophane radiated very little heat and mostly transmitted ambient thermal radiation reflected by the underlying aluminum foil. We can conclude that previous studies failed to show that ticks were attracted by radiant heat because the radiant heat stimulus used was much less intense than intended. Another flaw with this method is that if there had been a warm radiating surface in the arena, its radiation would have been reflected by the aluminum foil, confounding the location of the target. It is important that all surfaces in the arena are non-reflecting.

Several measurements and test conditions were used to determine whether the ticks in the thermotaxis arena were locating the warm TEC plate by radiant heat, warm air currents (convection) or heat conduction along the floor. Heating the TEC plates to 40°C increased the surface temperature of the floor less than 1°C and the air temperature less than 0.1°C, within 2.5 cm of the plate, with no observable change in temperature of either the plywood or air beyond that distance. To directly test the ability of ticks to detect radiant heat, we conducted trials with the TEC plate surfaces clad with aluminum foil (emissivity = 0.03) adhered with Glue Stic adhesive (Avery Dennison, Glendale, CA, USA). This would eliminate emission of radiant heat without affecting conductive or convective heat transfer to the ticks. We also conducted trials with IR-transparent 10 cm x 10 cm windows of polyvinylidene chloride film (PVDC, Saran™ food wrap) on wire frames placed 2.5cm from the TEC plates, to isolate air near the TEC plates from the rest of the arena and thereby eliminate possible convective but not radiant heat transfer to the tick.

To confirm that the Haller’s organ posterior capsule is the IR sense organ, thermotaxis trials were conducted with unfed virgin adult *A. americanum* females before and after application of low-melting dental wax (Kerr, Orange, CA, USA) onto the posterior capsules of both front legs using a hot 0.25 mm diameter silver wire attached to the tip of a soldering iron (Weller, Apex, NC). Care was taken to exclusively place the wax onto the posterior capsule; any wax accidentally applied to the anterior pit sensillae was removed using sterile forceps and a minute piece of kimwipe (Kimberly-Clark, Irving, TX). Following wax application, ticks were allowed to recover for 10 min prior to being tested in thermotaxis trials. We also attempted to localize the IR sense to the forelegs and the Haller’s organ by means of amputation, but the questing behavior of amputated test subjects was greatly reduced.

### Repellent Thermotaxis Bioassays

The effects of repellents on thermotaxis was assessed by introduction of known concentrations of repellent vapors into the closed air circulation system of the arena. An in-line low-pressure air circulation fan (Intex, New Delhi, India) pulled air from the arena through 1.5 cm-diameter screened exhaust ports, located midway up the side walls, 10 cm from each TEC plate, and re-introduced a combined airstream into the center of the arena through a screened 1.5 cm-diameter inlet port, at a rate of 900 ml/min (Fig. 1). Reagent grade repellents N, N-diethyl-meta-toluamine (DEET), 2-undecanone, picaridin, citronellal and nootkatone (Sigma Aldrich, St. Louis, MO, USA) were purchased 2 weeks prior to use and stored at room temperature. Repellents were diluted into 95% ethanol on the day of use so that 0.5 μl would give the desired concentration in the 10 l arena, which was applied to the tip of a temperature-controlled soldering iron and vaporized directly into the circulating air through an entry port in the suction tubing (Fig. 1) by briefly turning the tip temperature up to 65°C. Control repellent thermotaxis trials were performed with 0.5 μl 95% ethanol. Ticks were pre-exposed to the same control and repellent vapor concentration for 2 min in a 1l glass bottle prior to placement into the bioassay arena. Control and repellent thermotaxis trials used three runs of eight unfed virgin adult *A. americanum* females, unless otherwise noted.

### Y-tube olfactometer Bioassays

Y-tube olfactometer bioassays were conducted as previously described ^21^ to determine whether concentrations of repellents that disrupt thermotaxis also affect tick olfaction, another function ascribed to Haller’s organ. Tests were conducted during the day between the hours of 11:00-15:00, at 23 ± 1°C and 40% R. H., under ambient (LED) lighting. Each arm of the Y-tube olfactometer (Fig. S4) was supplied with either breathing quality air or 3% carbon dioxide (CO_2_) in breathing quality air (AirGas, NC, USA), at a flow rate regulated and measured with a #11 Compact Shielded flow meter (Gilmont Instruments, Barrington, IL, USA). A vacuum pump removed gases from the downwind end of the Y-tube at a rate equal to the total flow through the olfactometer and exhausted them out of the test area. Ticks were pre-exposed to repellent vapors for up to 5 min, as previously described, prior to placement into the Y-tube, with no additional repellent exposure during Y-tube bioassays. 50ng/cm^3^ and 100ng/cm^3^ DEET, which are 1X and 2X, respectively, the concentration found to completely abolish thermotaxis, were evaluated in Y-tube bioassays. After repellent pre-exposure, ticks were immediately placed at the Y-tube starting mark and their movements were filmed for 5 min. Olfactometer bioassays consisted of six runs of eight unfed virgin adult *A. americanum* females randomly selected from a total population of 250 and acclimated to experimental conditions for 20 min prior to repellent pre-exposure. To eliminate contamination and positional bias, all Y-tube components were washed thoroughly with 70% ethanol between replicates and the CO_2_ gas was alternated between the two arms of the Y-tube. Ticks were recorded as positive responders if they made a choice in Y-tube bioassays, moving 1 cm past the choice point (Fig. S4). Trials were filmed and scored as previously described and those data were analyzed using the Generalized Linear Model in R, as described above.

### Epi-Microscopy

The Haller’s organ posterior capsule was imaged with a Keyence VHX-5000 digital microscope using epi-illumination with polarized light and a VH-Z500R 500X-5000X objective (Keyence Corporation, Elmwood Park, NJ, USA). A Woodpecker UDS-J ultrasonic dental scaler (Guilin Woodpecker Medical Instrument Co., Guilin, China) was modified for micro-etching of the cuticle to remove the cover from the Haller’s organ posterior capsule for imaging. The stainless-steel tip supplied by the manufacturer was sharpened mechanically with a hard Arkansas whetstone wet with mineral oil, and then further electropointed in a solution containing 34 ml 98% sulfuric acid, 42 ml 25% orthphosphoric acid and 24 ml deionized water. A 6 VDC source was used with the variable resistance adjusted to control the rate of etching so that a sharp, polished tip could be obtained^24^. It was furthermore necessary to reduce the power supplied to the ultrasonic generator handpiece by placing a 3 KΩ, 5W resistor in series with it. The handpiece with sharpened tip were fixed to a micromanipulator to perform manually controlled etching of the cuticle covering of the Haller’s organ posterior capsule.

### Statistical Analysis

For scoring a thermotaxis assay as a choice test, ticks located within 10 cm of the warm plate (red area in Fig. 1A) at the end of the 5 min trial were counted as warm plate responders, while those ending up within 10 cm of the cold plate (blue area in Fig. 1A) were counted as cold plate responders. Ticks that did not reach either the warm or cold regions were counted as “neither” or “no choice”. Statistical analysis of choice data for each trial was performed using the generalized linear model ^25^ with binomial distribution and logit link function. Data were subjected to generalized linear mixed model ANOVA based on the model logit(Y_ij_) = U + T_i_ + e_ij_, using function glm in R3.3.3 (R Core Team 2017). In this model, Y_ij_ (Y=0,1) is the index of tick j (j=1,2,3… n) choice i (i=1,2,3), U is the average preference (probability) of tick j, T_i_ is the choice (preference) i effect, and e_ij_ is the residual error.

Estimated choice preference means were back-transformed to proportions. Pair-wise comparison results were given as response scale (i.e. difference of proportions, not log odds ratio). Significance of the overall preference effect was evaluated using a Chi-square test of deviance, and comparison of two choice preference proportion in terms of log odds ratio was conducted using a Wald test with z-values. P value adjustment: Tukey method for comparing a family of 3 estimates. A significance level of α = 0.05 (confidence level 95%) or better was used for all statistical tests.

---

**Figure S1.**
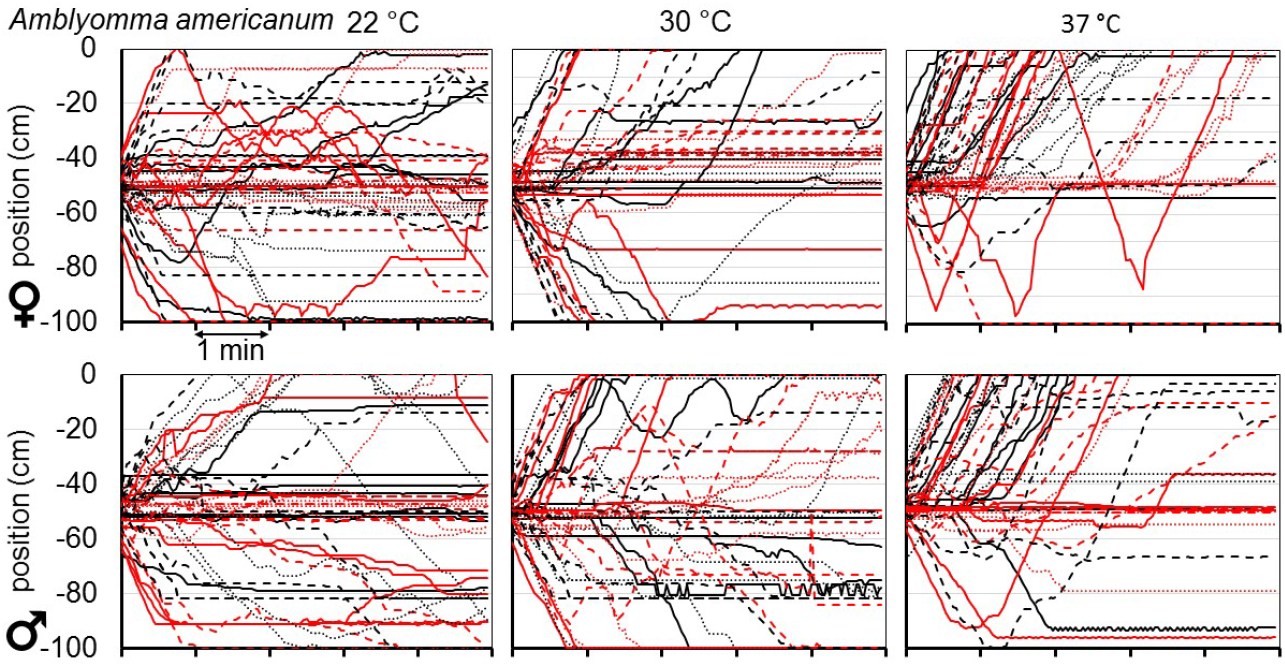
Longitudinal position vs. time in the 1m-long arena of 48 individual female (top) or male (bottom) *A. americanum* ticks (six runs of eight ticks, with a different color/line type combination for each run) after placement in the center, at −50 cm with the target plate (22, 30 and 37 °C in left, center and right panels, respectively) at 0 cm and the cold plate at −100 cm. Horizontal axis units are minutes.

**Figure S2.**
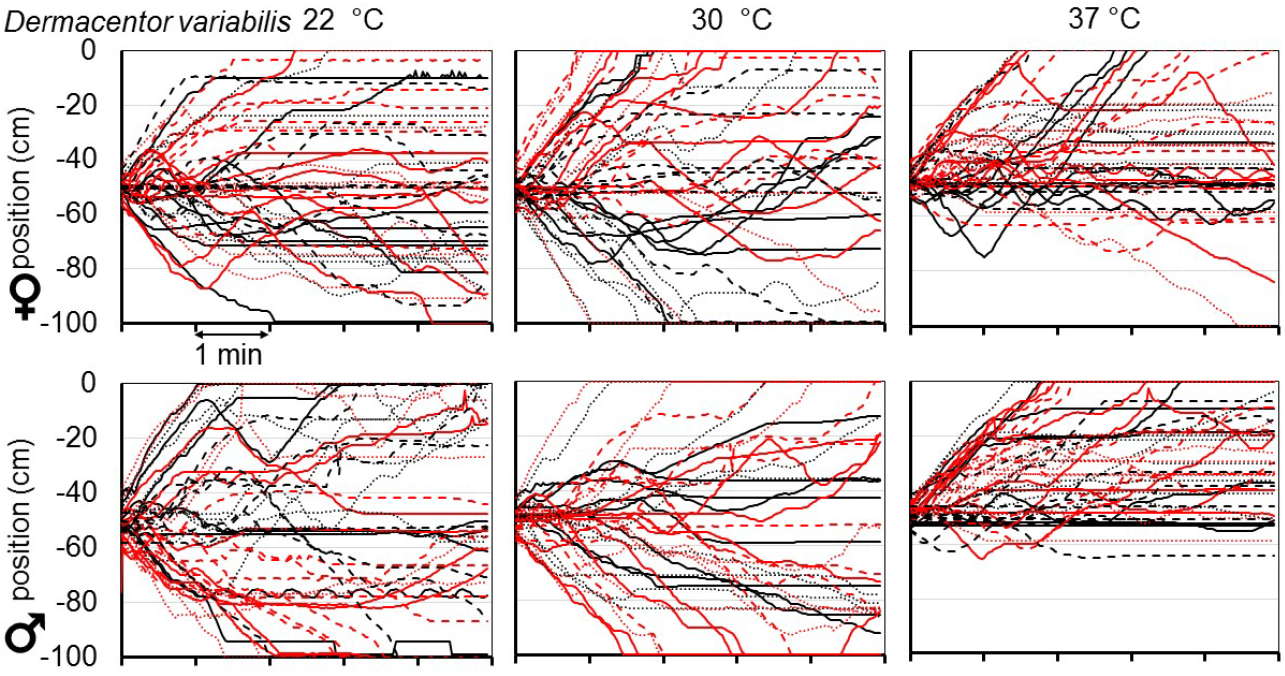
Thermotaxis trials for 48 individual *D. variabilis* female (top) or male (bottom) ticks, as in Fig. S1

**Figure S3.**
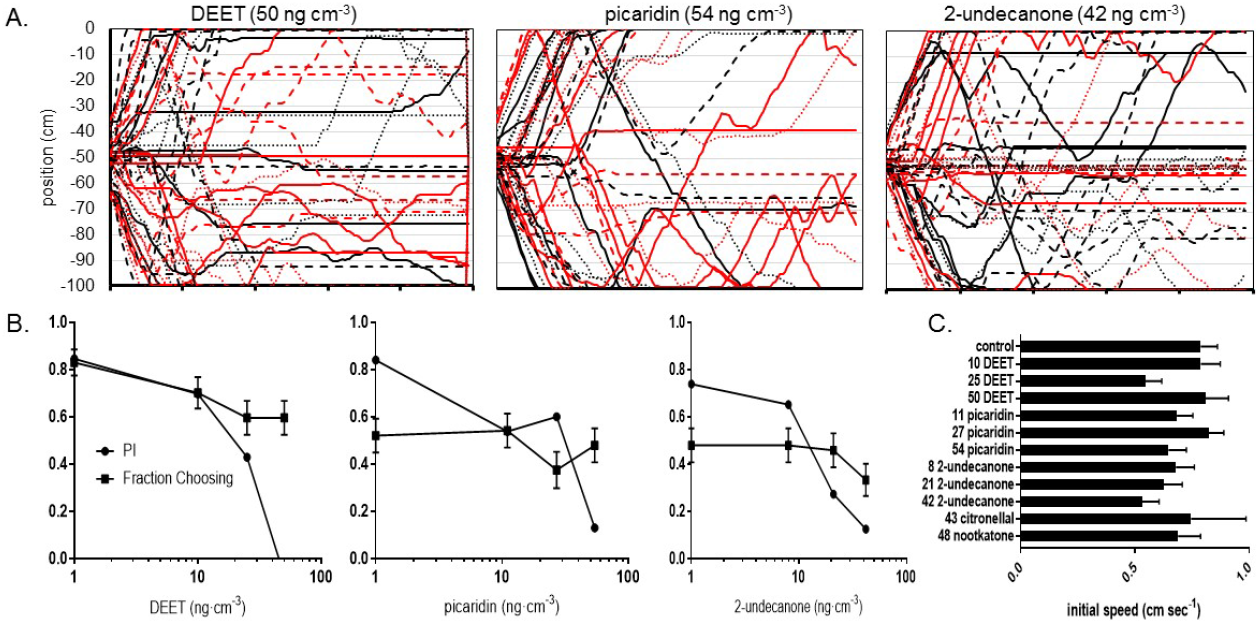
A. Each graph shows the longitudinal position vs. time of 48 individual *A. americanum* females (measured in six groups of eight, with a different line type for each group) after placement in the arena center, at −50 cm, as in Fig. S1, with the target plate at 40 °C and located at 0 cm and the room temperature plate at −100 cm. Horizontal axis units are minutes. Ticks were exposed to the indicated repellent at the indicated concentration before and during the thermotaxis trials. B. Concentration dependence of PI and fraction of ticks making a choice of either warm or cold, for DEET, picaridin and 2-undecanone, as indicated.

**Figure S4.**
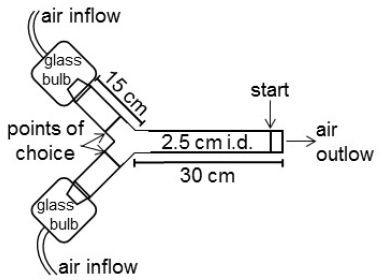
Y-tube olfactometer (from ^21^ with permission from Wiley & Sons, Inc.).

**Figure S5.**
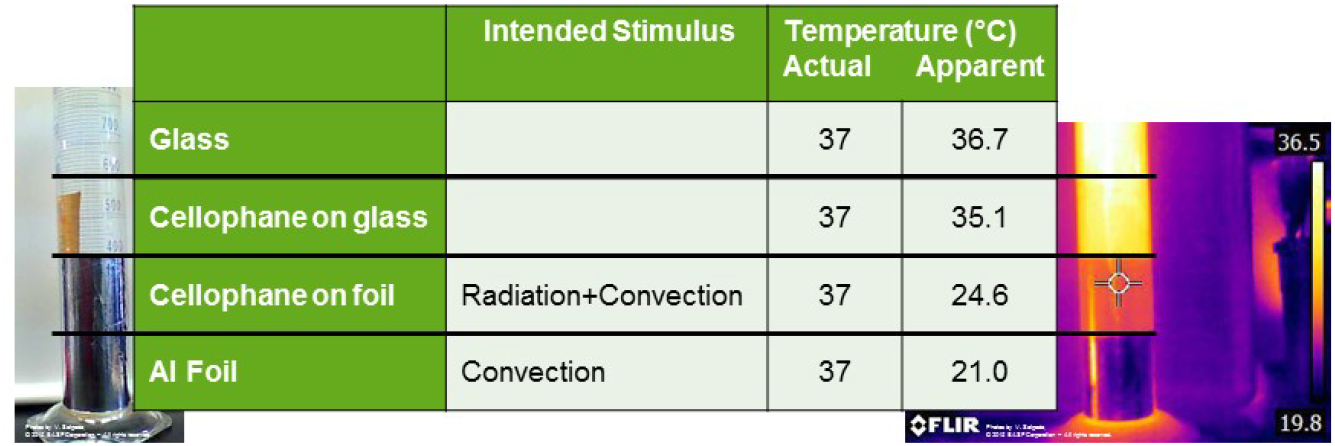
Demonstration that a thin, transparent layer of cellophane over aluminum foil, as used in previous attempts to demonstrate heat sensing by ticks (5) and human body lice (25) does not provide an effective radiant heat stimulus. A 1l graduated cylinder filled with water at 37 °C was wrapped partially with aluminum foil and partially with cellophane overlying the upper half of the aluminum foil, giving radiating surfaces of glass, cellophane over glass, cellophane over foil and foil alone. The visible light image is shown on the left and the thermal image is shown on the right, and the table shows the temperatures measured from the thermal image. The cellophane over aluminum foil section, which was intended in the cited studies to provide radiant heat, in fact radiated only enough heat to appear to the thermal imaging camera to be at 24.6 °C.

**Movie S1**. Three thermotaxis bioassay trials, each with 8 unfed virgin adult *A. americanum* females, demonstrating thermotaxis and its disruption by DEET. The air circulation tubes can be seen at the top of the arena and are positioned at −10, −50 and −90 cm. The TEC plates are just outside the left and right edges of the video. The dark areas on the left and right ends of the arena are shadows. The counters at each end show the number of ticks within 10 cm of the corresponding TEC plate and correspond to cold plate (left) or warm plate (right) choice scores. Top: Control with both plates at room temperature. Initially, four ticks go right and four go left. Final choice scores are 1 cold plate, 1 warm plate, 6 neither. Middle: warm plate at 37 °C. Five ticks go right initially and final choice scores: 0 cold plate, 5 warm plate, 3 neither. Bottom: warm plate at 40 °C in the presence of 50 ng/cm^3^ DEET. Initially, four ticks go right, three go left and one stays in middle initially but then moves left after a delay. Final choice scores are 1 cold plate, 1 warm plate and 6 neither.

## References and Notes

1. Wikel, S. K. Ticks and Tick-Borne Infections: Complex Ecology, Agents, and Host Interactions. Vet Sci 5, 60, doi:10.3390/vetsci5020060 (2018).

2. Sonenshine, D. E. & Roe, R. M. Biology of Ticks. Vol. II (Oxford University Press, 2014).

3. Carter, M. C. et al. Identification of alpha-gal sensitivity in patients with a diagnosis of idiopathic anaphylaxis. Allergy 73, 1131–1134, doi:10.1111/all.13366 (2018).

4. Lees, A. D. The Sensory Physiology of the Sheep Tick, *Ixodes Ricinus* L. Journal of Experimental Biology 25, 145–207 (1948).

5. Apanaskevich, D. A. & Oliver, J. H. J. in Biology of TIcks Vol. 1 (eds D. E. Sonenshine & R. M. Roe) Ch. 3, 59–73 (Oxford University Press, 2014).

6. Leonovich, S. A. Orientatinal Behavior of the Ixodid Tick Hyalomma Asiaticum Under Desert Conditions. Parazitologia 20, 431–440 (1986).

7. Carr, A. L. & Roe, M. Acarine attractants: Chemoreception, bioassay, chemistry and control. Pestic Biochem Physiol 131, 60–79, doi:10.1016/j.pestbp.2015.12.009 (2016).

8. Foelix, R. F. & Axtell, R. C. Ultrastructure of Haller’s organ in the tick Amblyomma americanum (L.). Zeitschrift fuer Zellforschung und Mikroskopische Anatomie 124, 275–292, doi:10.1007/BF00355031 (1972).

9. Steullet, P. & Guerin, P. M. Identification of vertebrate volatiles stimulating olfactory receptors on tarsus I of the tick *Amblyomma variegatum* Fabricius (Ixodidae). I. Receptors within the Haller’s organ capsule. J Comp Physiol A 174, 27–38 (1994).

10. Steullet, P. & Guerin, P. M. Perception of breath components by the tropical bont tick, *Amblyomma variegatum* Fabricius (Ixodidae). I. CO2-excited and CO2-inhibited receptors. J Comp Physiol A 170, 665–676 (1992).

11. Bruce, W. A. Posterior Capsule of Haller’s Organ in the Lone Star Tick, *Amblyomma americanum* (Acari: Ixodidae). The Florida Entomologist 54, 65–72, doi: 10.2307/3493790 (1971).

12. Materials. and methods are available as supplementary materials.

13. Costanzo, K. S., Schelble, S., Jerz, K. & Keenan, M. The effect of photoperiod on life history and blood-feeding activity in Aedes albopictus and Aedes aegypti (Diptera: Culicidae). Journal of Vector Ecology 40, 164–171, doi:doi:10.1111/jvec.12146 (2015).

14. Hardy, J. D. The Radiating Power of Human Skin in the Infra-Red. American Journal of Physiology-Legacy Content 127, 454–462, doi:10.1152/ajplegacy.1939.127.3.454 (1939).

15. Campbell, A. L., Naik, R. R., Sowards, L. & Stone, M. O. Biological infrared imaging and sensing. Micron 33, 211–225 (2002).

16. Mitchell, R. D., 3rd et al. Infrared light detection by the Haller’s organ of adult american dog ticks, Dermacentor variabilis (Ixodida: Ixodidae). Ticks Tick Borne Dis 8, 764–771, doi:10.1016/j.ttbdis.2017.06.001 (2017).

17. Bissinger, B. W. & Roe, R. M. Tick repellents: Past, present, and future. Pestic. Biochem. Physiol. 96, 63–79, doi:10.1016/j.pestbp.2009.09.010 (2010).

18. Sanford, J. L., Shields, V. D. C. & Dickens, J. C. Gustatory receptor neuron responds to DEET and other insect repellents in the yellow-fever mosquito, Aedes aegypti. Naturwissenschaften 100, 269–273, doi:10.1007/s00114-013-1021-x (2013).

19. Dickens, J. C. & Bohbot, J. D. Mini review: Mode of action of mosquito repellents. Pesticide Biochemistry and Physiology 106, 149–155, doi:10.1016/j.pestbp.2013.02.006 (2013).

20. Sokabe, T., Chen, H. C., Luo, J. & Montell, C. A Switch in Thermal Preference in Drosophila Larvae Depends on Multiple Rhodopsins. Cell Rep 17, 336–344, doi:10.1016/j.celrep.2016.09.028 (2016).

21. Carr, A. L. et al. Responses of Amblyomma americanum and Dermacentor variabilis to odorants that attract haematophagous insects. Med Vet Entomol 27, 86–95, doi:10.1111/j.1365-2915.2012.01024.x (2013).

22. Schneider, C. A., Rasband, W. S. & Eliceiri, K. W. NIH Image to ImageJ: 25 years of image analysis. Nat Methods 9, 671–675 (2012).

23. Wigglesworth, V. B. The sensory physiology of the human louse Pediculus humanus corporis de Geer (Anoplura). Parasitology 33, 67–109 (1941).

24. Grundfest, H., Sengstaken, R. W. & Oettinger, W. H. Stainless Steel Micro-Needle Electrodes Made by Electrolytic Pointing. Review of Scientific Instruments 21, 2, doi:https://doi.org/10.1063/1.1745583 (1950).

25. McCullagh, P. & Nelder, J. A. Generalized linear models. 2nd edn, (Chapman and Hall, 1989).

